# Time-controlled Multichannel Dynamic Traction Imaging of Biaxially Stretched Adherent Cells

**DOI:** 10.1101/2020.03.02.972919

**Authors:** Aron N. Horvath, Andreas A. Ziegler, Stephan Gerhard, Claude N. Holenstein, Benjamin Beyeler, Jess G. Snedeker, Unai Silvan

**Affiliations:** Department of Orthopedics, Balgrist University Hospital, Zurich, Switzerland; Department of Health Sciences and Technology, Institute for Biomechanics, ETH Zurich, Zurich, Switzerland

**Keywords:** dynamic TFM, cellular relaxation, nuclear deformation, mechano-active substrate, strain rate

## Abstract

Here, a dynamic traction force microscopy method is described which enables sub-second temporal resolution imaging of transient subcellular events secondary to extrinsic stretch of adherent single cells. The system employs a novel tracking approach with minimal computational overhead to compensate substrate-based stretch-induced motion/drift of stretched single cells in real time, allowing capture of biophysical phenomena on multiple channels by fluorescent multichannel imaging on a single camera, thus avoiding the need for beam splitting with associated loss of light. The potential impact of the technique is demonstrated by characterizing transient subcellular forces and corresponding nuclear deformations in equibiaxial stretching experiments, uncovering a high frequency strain-rate dependent response in the transfer of substrate strains to the nucleus.

---

Understanding how changes in the mechanical environment alter cellular functions is a central goal of mechanobiology. Several approaches have been developed to mechanically challenge cells in a controlled fashion and to uncover subsequent adaptation with detailed focus on magnitude, frequency and duration of the external stimuli.^[1–4]^ However, not all methods allow simultaneous imaging-based quantification of the cellular response. Shear flow,^[5]^ optical tweezer,^[6]^ AFM and micropipette aspiration techniques induce negligible sample-motion during mechanical perturbation,^[7.8]^ however the large displacements and high strain rates inherent to tensile testing make the tracking and imaging of the stretched cells especially demanding. ^[9]^

Previous studies in which cells were mechanically tested in tension relied on manual compensation of the drift.^[10–13]^ However, none of the existing approaches can automatically compensate the motion of the stretched cells and record the cellular response without substantial time delay and in a time-controlled manner. Such a method would allow the investigation of spatiotemporal involvement and role of different cellular components in stretch evoked cellular behavior.

Here we present a single camera-based tracking microscope capable of real-time counteraction of motion of the stretched sample in three dimensions and acquiring multichannel fluorescence images for quantification of the resulting cellular responses. The system is based on a frame of a widefield upright microscope (Leica DM5500) equipped with a 40X water immersion objective, a CCD camera and a multichannel LED light source that can capture several fluorescently stained cell compartments of the mechanically challenged cells (**Figure 1a**). The substrate deformation was controlled by a pressure-actuated mechanical stretching device delivering equibiaxial strain to the cells seeded onto (silicone-membrane supported) stretchable polyacrylamide (PAA) hydrogel (**Figure S1 and S5**). To resolve the cellular traction forces and to determine the surface strain, fluorescent beads encapsulated close to the substrate surface were used as fiducial markers. The compensation of the sample-motion was managed by a motorized stage that utilizes a novel computer vision algorithm described below.

**Figure 1.**
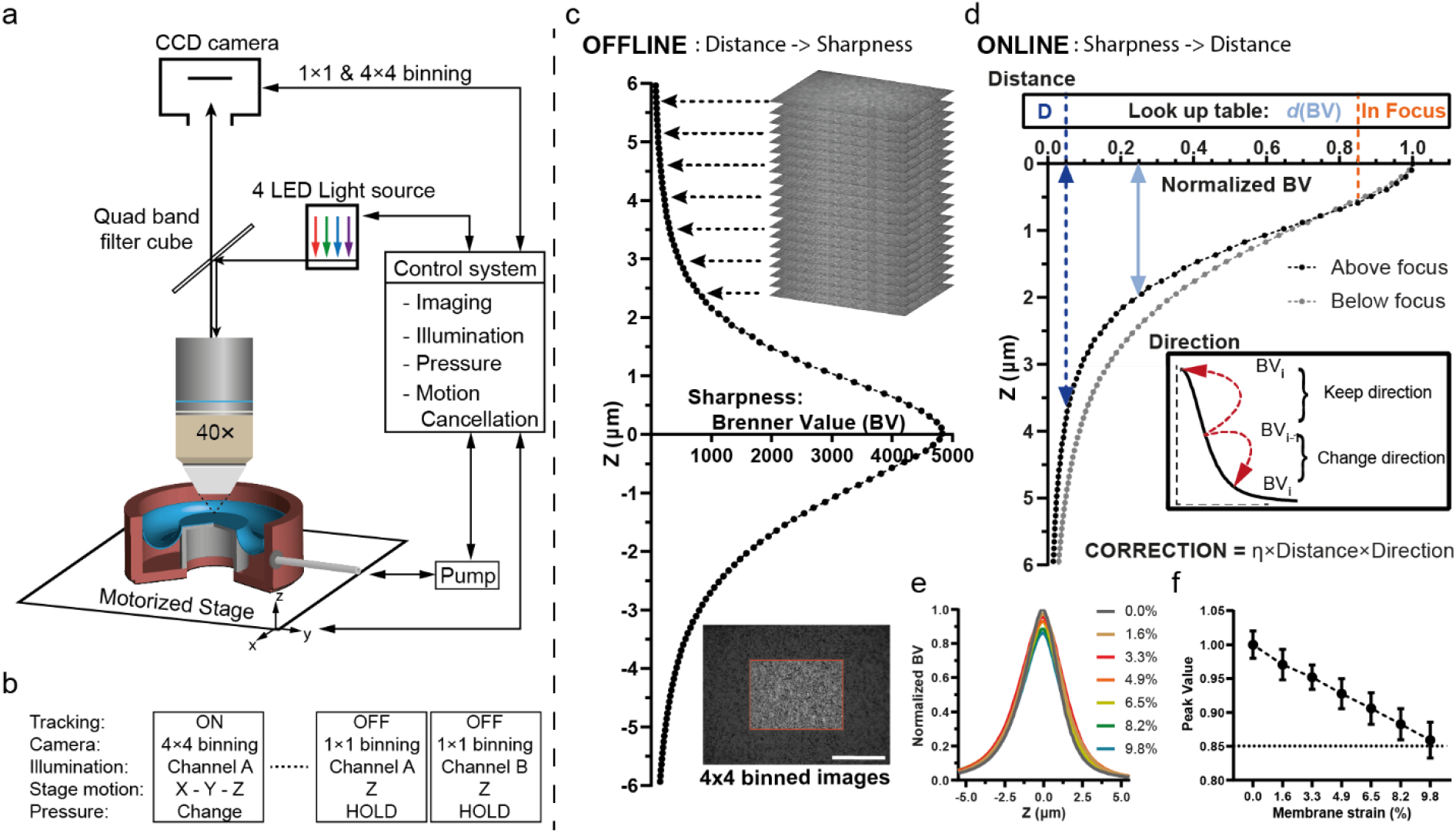
Tracking microscope for imaging of mechanically stimulated cells. (**a**) Schematic illustration of the system, which consists of a widefield fluorescence microscope equipped with a motorized stage, 4 channel LED light source and a pressure-controlled vacuum-actuated equibiaxial stretching device (**b**) The camera is used in 4×4 binning mode for motion-cancellation by extracting the lateral and axial drift of the sample from the identically-illuminated images and compensating them by the motorized stage. During high-resolution imaging the tracking is suspended, and the camera is set to 1×1 binning mode. To capture multiple compartments of the cell Z-stacks are acquired sequentially by sequentially changing the light channels. (**c-d**) Focus tracking is relying on sample-specific information acquired before stretching (offline) and used during the tracking procedure (online). (**c**) The offline tracking algorithm consists in the acquisition of a Z-stack (4×4 binned images) around the focal plane and in extraction of image-contrast based sharpness using the widely applied Brenner function for each image center to create a sample specific distance-sharpness curve (**d**) The online tracking algorithm uses the distance-sharpness curve in an inverse fashion to estimate the distance from the ideal focus position based on the calculated image sharpness. Two threshold values are introduced to stabilize the tracking performance: (1) a lower one to minimize correction error stems from large distance difference between small sharpness values, below this value the distance of the correction is kept constant equivalent to the correction value of the threshold sharpness itself (D); (2) above the upper threshold the sample considered to be in focus to compensate the error in distance estimation caused by the lateral drift of the sample. The direction of the applied stage correction is determined based on the previous sharpness value. The correction was adjusted with a damping parameter η set between 0.9 and 1. (**e**) Robustness of sharpness-distance curve over pressure change was measured by acquiring distance-sharpness curves at different membrane strains. An example for the change of the sharpness-distance curve over strain values ranging from 0 to 10% is shown. Although the maximal sharpness is decreasing, the overall characteristic of the distance-sharpness curve is not affected. (**f**) The peak sharpness value is linearly decreasing with the membrane strain, so the upper threshold was adjusted according to the maximum aimed membrane strain. (n = 4 membranes; 9-12 positions per membrane were analyzed). Scale bar (c): 100 μm

The algorithm separately determines and cancels the lateral and axial drifts of the sample by adjusting the stage-position relative to the actual one. For tracking, we used the camera in a 4×4 binning mode with a resolution of 348 x 260 pixels providing 70 fps. Besides the advantage of the high imaging frequency, the binned images require less computational power and decreased light intensity resulting in significantly reduced phototoxicity. While the lateral drift is determined by calculating relative spatial shift between the subsequently acquired real-time images using fast Fourier transformation (FFT) – based digital image correlation, the axial motion-cancellation relies on a newly developed autofocus method based on image contrast.

This autofocus algorithm is composed of two parts: (1) an offline step to generate information of the focus position relative to the surrounding focus planes for (2) the online, supervised mountain-climbing search algorithm which is applied to auto-focusing. In the offline step (**Figure 1b**), the algorithm performs a Z-axis sweep with 100 nm steps around the focus of the sample-specific surface region and extracts an image contrast-based sharpness value for each plane through the Brenner function (used in autofocusing approaches) using only the center (174 × 130 pixels) of the binned images.^[14,15]^ Considering the position of the desired focus plane as reference, a relation between the distance from the desired focus plane and the Brenner (sharpness) value [BV] can be defined. The new algorithm assumes that the distance-sharpness curve is symmetric around the focus plane with the maximum BV, which is true for the surface-aligned beads. Importantly, any tracking feature which meets the stated criteria, such as stained nuclei, is suitable for tracking.

Using this curve in an inverse fashion, the online auto-focus matches the extracted BV of the real time acquired images (< 1ms) to the corresponding distance value. The focusing method only relies on the narrower side of distance-sharpness curve to avoid overestimation of the distance (**Figure 1c**). As the axial and lateral drift-compensations occur simultaneously, the BV of the acquired images is not only affected by the camera noise, but potentially also by the error in the lateral motion-cancellation. To test whether the lateral drift affects the focusing performance, we recorded several distance-sharpness curves at different positions of the membrane. Thereof we determine and introduce a lower and upper threshold as well as an additional damping parameter (η) for stabilization. The distance value for the BVs below the lower threshold are maximized by the distance value (D) corresponding to the threshold as the difference in small BVs are resulting in very large distance differences. However, above the introduced lower threshold the error in the correction is less than +−0.5μm, which is below the theoretical depth of field of the microscope (**Figure S3**). Thus, images with required equal or smaller distance adjustment (based on their BVs) are considered to be in focus. The direction of the correction is initialized according to direction in which the axial drift is expected, but it is constantly updated by comparing the sharpness of the current and the previous image (**Figure S2**). To confirm that the same sharpness-distance curve can be used during the stretching experiment, BV curves were acquired at different stretched states (**Figure 1e**). The maximum of the curve was linearly decreasing with increased membrane strain; however, the overall characteristic of the curve did not change (**Figure 1f**). Considering this, the upper threshold was adjusted according to maximal aimed strain value. To minimize lateral tracking error, we used a closed loop controller implemented with a Smith Predictor to manage the time delays caused by camera, stage and offset calculation, for the lateral compensation.^[15]^ For the axial compensation, images were only considered after each finished focus adjustment.

We applied the tracking microscope to assess real-time changes of the cell contractility and nuclear deformation as a response to applied strain-rate-dependent tension. Cells seeded on PAA hydrogels with 45×45 μm^2^ fibronectin micropatterns and incorporated fluorescent markers were exposed to 5% equibiaxial static strain at two different strain rates: 0.5%/s or 5%/s (**Figure 2a,f,h**). To capture the cellular response, asynchronous multichannel imaging was performed with the same camera used for tracking but without pixel binning at 20 fps. To allow postprocessing compensation of any potential tilt of the sample, a sequential Z-stack was acquired for each channel turning on/off each LED light source between images. In the experiments we describe here, bead displacement and nuclear projected area were quantified before and after substrate deformation within the first 75 seconds (**Figure 2b,c**), with the long-term behavior of the cells being recorded for at least 25 minutes. To the best of our knowledge, this was the first time that the relaxation of cells exposed to static stretch could be quantified in a time-controlled manner at sub-second resolution.

**Figure 2.**
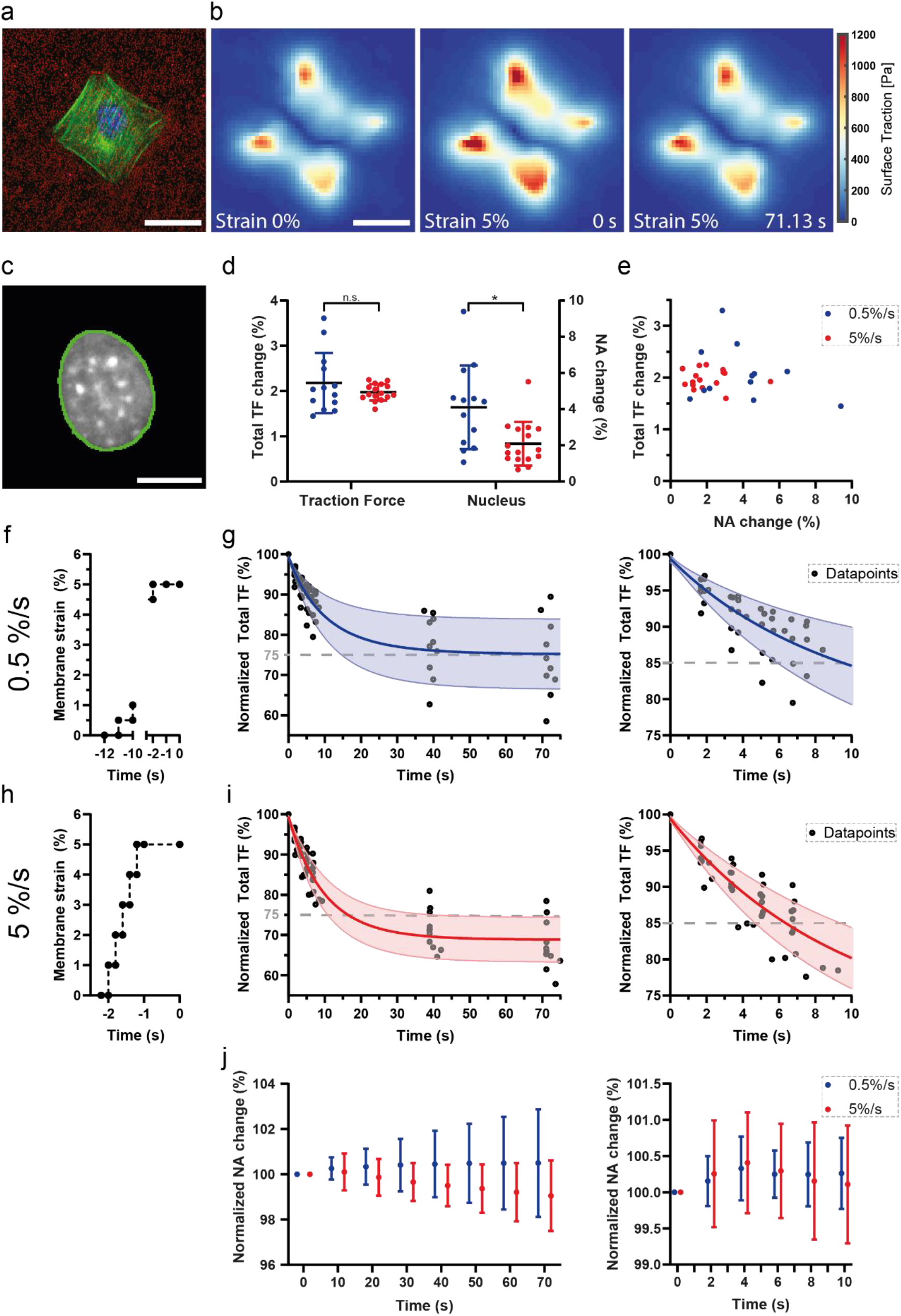
The tracking microscope enables accurate observation of changes in traction force and in nuclear projected area after 5% substrate stretch in a time-controlled manner. (**a**) Example of a 3T3 fibroblast adhered to a 45×45 μm2 fibronectin-coated pattern (red - fluorescent labelled microspheres; green – actin network, blue - nucleus), Scale bar 25 um. (**b**) Example for quantified traction force heatmaps showing rapid cell relaxation after the initial traction force changes: non-stretched (left panel), initial at 5% strain after 5%/s strain stretch (middle panel), and after 71.13 s at 5% strain (right panel). (**c)** Example for nuclear area segmentation (green line – segmented perimeter of the nucleus). (**d**) Strain-rate dependent transient response of the cellular contraction and nuclear deformation was quantified relative to the baseline. Average nuclear area (NucA) and averaged total traction force (TF) values for the two stretching conditions (0.5%/s, TF: 2.18 +/− 0.66 fold, NucA: 4.11 +/− 2.31% n = 13 cells; and 5%/s, TF: 1.97 +/− 0.18 fold, NucA: 2.09 +/−1.21% n = 16 cells), (**e**) No correlation found between change in (NucA) vs change in average total (TF) of either 0.5%/s or 5%/s stretched cells. (**f,h**) Stretching profile of the performed 0.5%/s (step size: 0.5%, time interval = 1 second) and 5%/s stretch (step size: 1%; time interval = 0.2 second) with an extended 1 second delay. (**g,i**) Total (TF) relative to initial (TF) (at 5% t = 0 s) within the first 75 second (left panels) or 10 second (right panels), the average relaxation curve was obtained by averaging the individually fitted exponential curves on the corresponding datapoints (dot - individual datapoints, solid line – average fitted value, transparent area – standard deviation). (**j**) Relative nuclear area changes within the first 75 second (left panel) or 10 second (right panel), the mean values with standard deviation were obtained by averaging the individual interpolated nuclear values. Scale bar: (**a,b**): 25 μm (c): 10μm; Statistic (**d**): n.s., not significant, *, p<0.01.

The average transient change of the total traction force and of the nuclear area were strain-rate-dependent showing smaller changes at the higher strain rate (**Figure 2d**). Specifically, the change in traction force and nuclear area were 2.18 +/− 0.66 fold and 4.11% +/− 2.31 respectively when stretched at 0.5%/s and 1.97 +/− 0.18 fold and 2.09% +/−1.21 at 5 %/s). However, there was no correlation between the two quantified cellular responses (**Figure 2e)**.

The traction force relaxation and nuclear area change during the static 5% strain relative to the first (t=0s) time point was obtained by fitting an exponential curve and interpolating between the individually captured datapoints for each analyzed cell. We found that the average total cellular traction was dramatically decreased in a strain-rate dependent manner within the first seconds after the application of the tensile stress. Specifically, cells stretched at 0.5%/s relaxed their contractility by 15% in the first 10 and by 25% in the first 70 seconds compared to those stretched at 5%/s in which the same relaxation was reached after 6 and 16 seconds, respectively (**Figure 2g,i, Figure S4**). In case of the nucleus, the average change compared to the initial state was not significant for any of the tested stretching speeds (**Figure 2j**).

In conclusion – we have implemented a novel high-speed tracking microscope with sequential multichannel imaging for the examination of stretch-induced changes in cell contractility and deformation of the nucleus. We demonstrated that the system can be used to investigate transient (sub-second) cellular responses, which have been previously inaccessible in similar stretching experiments. The quantified changes of cellular contraction in static strain relaxes exponentially which strongly underlines the necessity of accurate temporal observation. The hereby quantified changes in traction forces relative to baseline not only extend recently published results,^[12]^ but confirm the strain-rate-dependency reported from stiffness-clamp experiments.^[17]^ In apparent lockstep with transient cellular contractility, recorded nuclear deformations followed the observed strain-rate dependent behavior indicating transiently elastic coupling between the ECM and nucleus,^[18,19]^ but reflected the longer term viscoelastic relaxation that has been reported by others.^[20]^ With this we demonstrate how the presented system opens up the possibility to uncover the mechanical connectivity and time-dependent engagement of different cellular compartments in response to tensile perturbations of different natures.

Our results also demonstrate the applicability of image-contrast-based methods for real-time tracking purposes. The sample-specific information-extended algorithm eliminates the iteration extensive (mountain climbing) focus search by a precise estimation of required stage correction with accuracy of a few hundred nm.^[15]^ The generality of such focusing algorithm enables it to be used with any fast fluorescence microscope technique where the Brenner function-based interpretation of sample sharpness gives quasi-symmetric sharpness-distance curve around the desired focus plane.

In alternative embodiments, the tracking microscope could potentially be extended to a multi-camera-based system with a separately dedicating a camera for low-resolution tracking and one for simultaneous multichannel high-resolution imaging, allowing continuous motion-cancellation parallel to high resolution image acquisition.

## Experimental Section

### Hardware for lateral and axial motion cancellation

The motion cancellation system described here is built around a Leica DM5500 upright microscope equipped with Leica DFC360 FX monochrome digital camera and a HCX APO 40X/0.80 water immersion objective. The lateral-motion cancellation utilized an automated xy motorized stage (Marzhauser), and the axial-movement was performed by the Leica Z-drive. The stage was driven and controlled in all three dimensions by a PCI card (Oasis blue, Objective imaging) installed on a personal computer (PC) as a part of the custom developed tracking system.

### Stretching device

A commercially available stretching device (Flexcell StageFlexer) was used to apply equibiaxial strain to the silicone membrane. A vacuum pump (V-700, Büchi) evacuates the small chamber of the device resulting a pressure difference leading to deformation of the round silicone membrane (SMI silicone sheet, thickness 0.02 inch) by drawing it over a loading post. To precisely regulate the pressure, a pressure controller (T 3110, Marsh Bellofram) was used. The controller consisted of two valves: one connected the chamber to the pump and the other one the chamber to the environment. By closing and opening of the valves, the actual chamber pressure was regulated. The communication between the PC and the pressure controller was through a digital/analogue converter (NI 6009, National Instruments) allowing to set and monitor the pressure within the chamber. To match the converter’s output range to the controller’s set point, the communication was extended with a custom-built voltage converter. Due to the nature of the pressure controller, the platform was functioning in a stepwise/staircase fashion (**Figure S5**). The desired target pressure was divided into defined number of steps with defined time intervals. We applied a silicone oil as lubrication between the membrane and the cylindric support stage with a viscosity of 1 Pa·sec.

### Membrane Calibration

To calibrate the stretching of the hydrogel (polymerized on top of the silicone membrane) relative to the applied vacuum-pressure, the substrate was stretched and imaged within the actuator-pressure range [0 60kPa]. The surface was determined based on the displacement of the embedded fluorescent beads showing linear behavior (**Figure S5**).

### Focus tracking algorithm

To generate the sharpness-distance curve around the focus plane during the offline part of the algorithm, 5 images were recorded at each Z-stage position and the corresponding BVs were averaged. This allowed suppression of potential errors rising from imaging noise. The introduced lower threshold stabilizes the focusing performance by maximizing the distance value for the BVs below the threshold (Fig. 2d) as the difference in small BVs are resulting in very large distance differences. The lateral drift caused inaccuracy in distance estimation was compensated by the introduced upper threshold. The damping parameter (η) was set between 0.9 and 1 as an additional stabilization.

### Light source

The CoolLED pE-4000 illumination system was controlled remotely by a custom developed PCB board and a microcontroller (Arduino Uno R3) via 4 TTL inputs coupled with 4 analog inputs for independent on/off and intensity control of each light channel. The PCB board comprised of a 12-bit digital-analog converter (Adafruit, MCP4725 Breakout Board), an operational amplifier (Texas Instrument, LM358PE3) and a multiplexer with four differential channels (Analog Devices, ADG409). The communication between the PC and microcontroller consisted of a serial PCI express card (Delock 89333) and RS232 Arduino Shield (Seeed Studio) with an average response time <1ms.

### Sample preparation

Stretchable, fibronectin micropatterned polyacrylamide hydrogels with surface aligned beads were prepared according to previously published protocol (**Figure S1**)^[21]^ Briefly, array of 45×45 μm2 square fibronectin micropatterns were generated by standard microcontact printing technique on an ethanol and water washed coverslip. Then, silicone membranes (diameter 47 mm, thickness 0.5 mm) were incubated benzophenone (10 % w/v in 35/65 w/w water/acetone) adsorption for 1 minute then washed with methanol twice. The membranes were placed in a desiccator for 30 minutes then flushed with N2. In the meantime, polyacrylamide prepolymer were prepared by mixing acrylamide/bisacrlyamide/acrylic-NHS (10/0.13/0.005/ w/v), and 1 M HCL/tetramethylethylenediamine/0.5 μm diameter green fluorescent beads (0.0054/0.0005%/0.02% v/v), respectively. The polymerization was initiated by adding ammonium persulfate (0.02% v/v) to the solution. The benzophenone adsorbed membrane and the micropatterned coverslips with a 0.18 mm spacer were assembled into a sandwich filled with the polymerizing the prepolymer. This construct was centrifuged at 4200 g for 15 minutes in order to align the beads to the hydrogel surface. Then the coupling between the hydrogel and the membrane was initiated by 10 minutes deep UV activation of the adsorbed benzophenone. After complete polymerization, the samples glass coverslip was separated under PBS then hydrogel surface was passivated with 5% BSA in PBS for 1 hour. After the incubation, the samples stored at 4°C in PBS prior use.

### Cell culture and staining

NIH/3T3 fibroblast were cultured at 37°C, 5% CO2, in DMEM F12 (FluoroBrite™ DMEM, A1896701), supplemented with 10% FBS and with 1% Penicillin/Streptomycin. Cells after accutase solution-based dissociation (Sigma, A6964) were seeded at 81 cells/ mm2 on the hydrogels and allowed to adhere to the patterns for 12 h. Two hours before the experiment, the cells were stained by the addition of the live-dye SiR-actin (SpiroChrome), Hoechst 33342 (NucBlue™, Live ReadyProbes™ Reagent, R37605) and anti-fade solution (ProLong™ Live Antifade Reagent, P36974) at a final concentration of 500nM, 5%v/v and, 2%v/v, respectively.

### Fluorescent image analysis

The full-frame imaging with resolution of 1392×1040 pixels were performed at 20 Hz. For each fluorescence channel, an image-stack was acquired with 0.7 μm intervals for a total distance of 7 μm (if required up to 10μm) to compensate the tilt of the sample. For the observation of force- and nuclear-relaxation, Z-stacks were acquired at high frequency within the first 10 seconds followed by lowering imaging frequency. The traction forces were estimated by quantification of the surface deformations from the acquired Z-stacks of the fluorescent beads. The acquired images were aligned using FFT-based cross correlation to eliminate any shift within the recorded Z-stacks. To compensate the tilt of the sample, maximal intensity projection was applied to the Z-stacks after removing the highly blurred images based on their sub-regional sharpness assessed by the Brenner function. To determine surface traction, samples were treated with 0.1% SDS (0.1% in PBS). As the lubrication failed after 30 minutes static strain, reference image at 0% strain was created by translation and affine transformation of the reference image at 5%.^[12]^ The quantification of cell tractions was performed as previously described.^[22]^ The nuclear area was determined by standard segmentation method from the sharpest nuclear image within the acquired Z-stack. These discrete set of data were used to derive expected continuous-changes of the quantified changes by fitting exponential decay curves to each contractility relaxation and by interpolating between each nuclear datapoints using shape-preserving piecewise cubic interpolating polynomial. The analysis of the transient cellular response was extended with additional data from only transient response measurements.

### Statistical analysis

Statistics and sample size are reported in the figure legends and text for each measurement. The different strain-rate-dependent cell contractility and nuclear projected area were compared by unpaired T-test (each dataset was normally distributed tested by Kolmogorov-Smirnov test with Lilliefors Significance correction)

### Data availability

The data that support the findings of this study are available from the corresponding authors upon request.

### Code availability

Software for imaging and microscope control was written in C++, using the QT framework and other third-party libraries. Image and data processing were implemented in MATLAB R2016b. Software for operating the tracking microscope and for image/data processing, as well as license information and implementation notes, are available upon request.

## Supporting Information

Supporting Information is available from the Wiley Online Library or from the author.

## Acknowledgements

We thank Tobias Klauser for help with software development, especially involvement in the design of the tracking algorithms. This work was supported by the Swiss National Science Foundation (grant numbers 165670, 138221, 118036 to J.G.S.)

## Supporting Information

**Figure S1.**
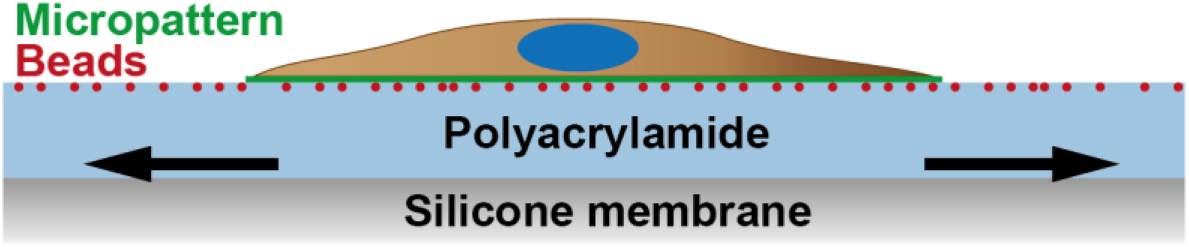
Schematic illustration of biaxially stretched substrate. A fibronectin micropatterned polyacrylamide hydrogel was polymerized onto a silicone membrane as a support substrate. The fluorescence beads were aligned to the substrate surface by centrifugation before polymerization.

**Figure S2.**
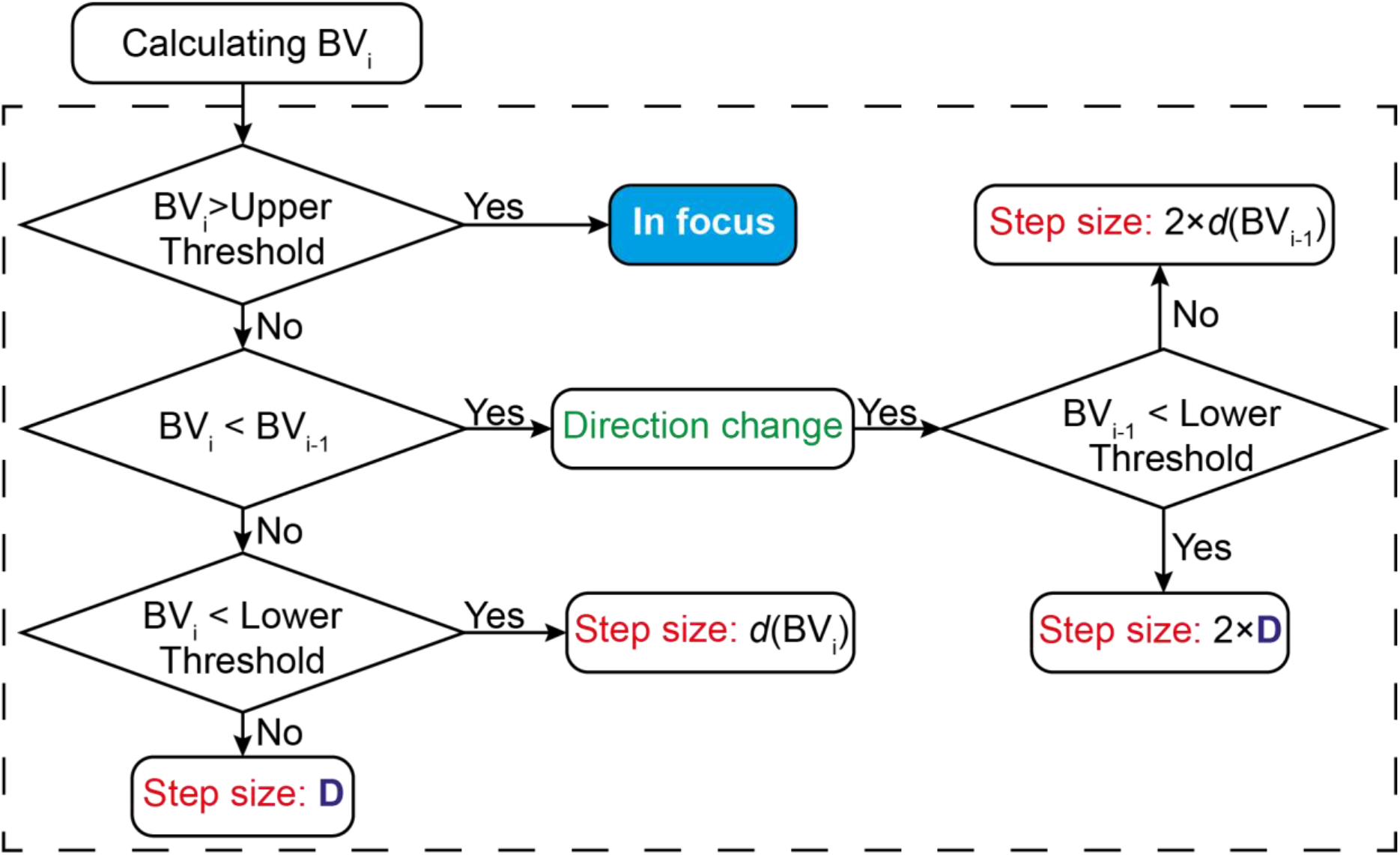
Flow chart of the online tracking algorithm. First, the algorithm calculates the BV_i_ for the captured image I_i_ and compares the defined upper threshold value. If BV_i_ is larger than the threshold, the sample is considered to be in focus and no correction is performed; if smaller then it further compares to the value of the previously captured image (BV_i_-1). In case of BV_i_ is larger than BV_i_-1), the BV_i_ is compared to the lower threshold to define the size of the correction step. If it is larger than the lower threshold than the correction value is determined based on the sharpness-distance curve (*d*(BV_i_)); otherwise the step size corresponding to the threshold set (*D*). If BV_i_ was smaller than the BV_i_-1 (meaning that the previous correction was performed in the wrong direction), the direction of the following correction is reversed, and the step size is doubled.

**Figure S3.**
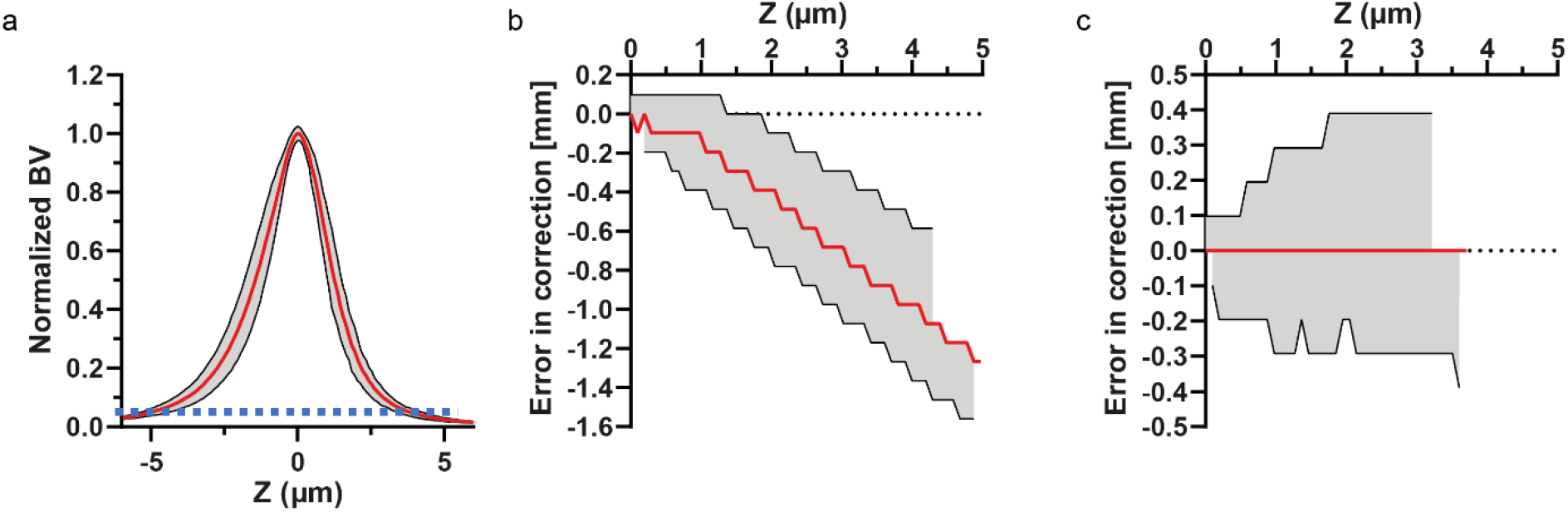
Quantitative evaluation of the Brenner function-based sharpness distance curves between different positions of the substrate surface. (**a**) Several sharpness-distance curves were acquired at different locations of the same sample to test the potential effect of the lateral shift on the focusing performance. Normalized average of sharpness-distance curves is shown by the red line, the gray area indicates the standard deviation and the blue dashed line represents the lower threshold with 0.05, (n = 9 positions) (**b-c**) Taking the average of the curves, we determined the error in correction within the observed standard deviation. The error was calculated considering both sides of the curve for BV equal or above the 5% of the maximal BV (lower threshold) by using the narrower (right) side of the average curve to determine the intended correction step size. (**b**) The extracted BV for images below the desired focal plane (left side of the curve) resulted increasing under-estimation of the steps with increasing distance from the peak. (**c**) For BVs above the focus plane, the observed difference within the recorded deviation is below 0.4μm.

**Figure S4.**
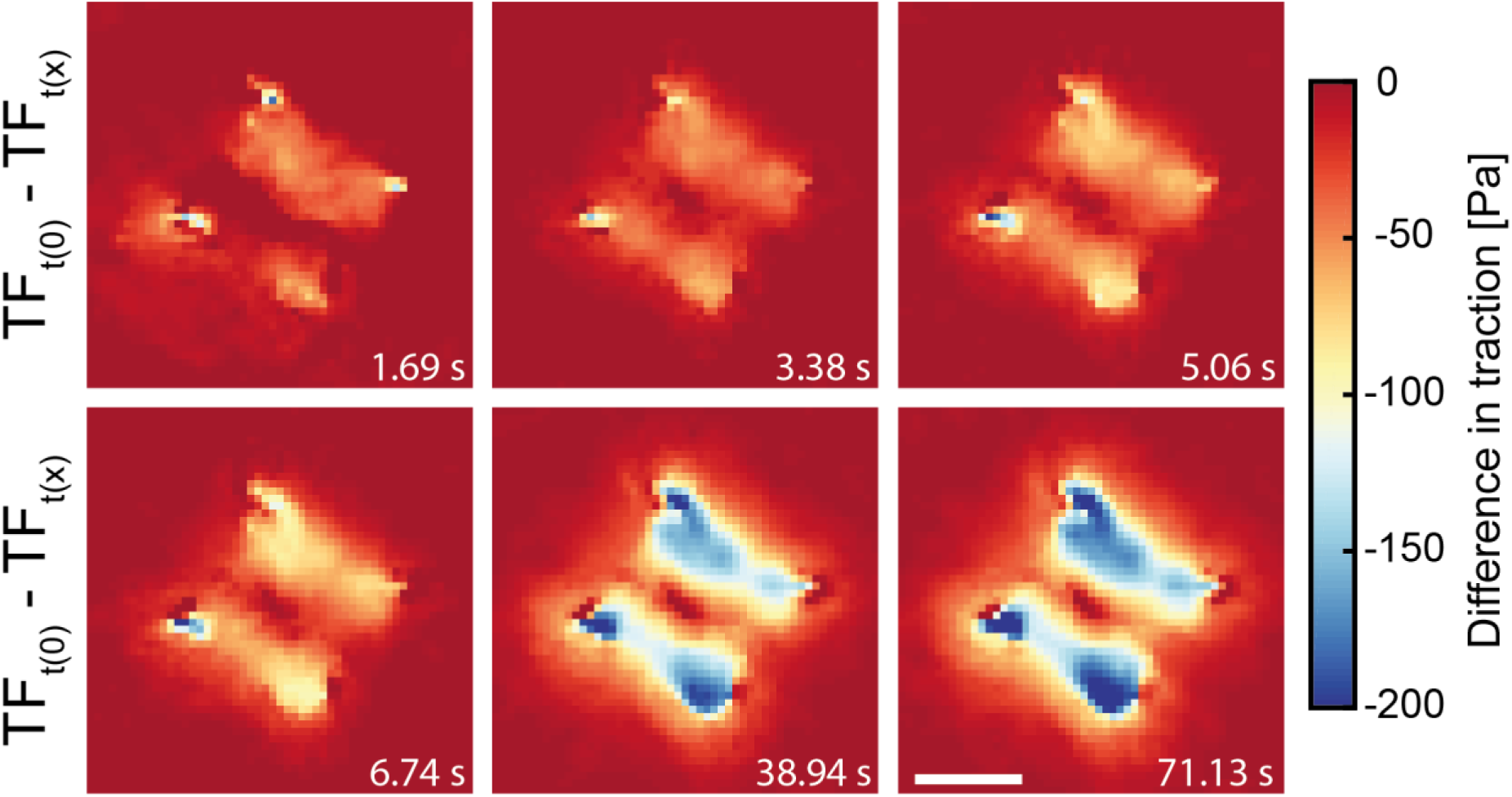
An example of cellular relaxation. The surface traction generated by a 3T3 fibroblast exposed to 5% stretch with 5%/s strain rate decreases rapidly relative the initial state of the cell (considered to be at t = 0 s) illustrated by the traction (heat) maps.

**Figure S5.**
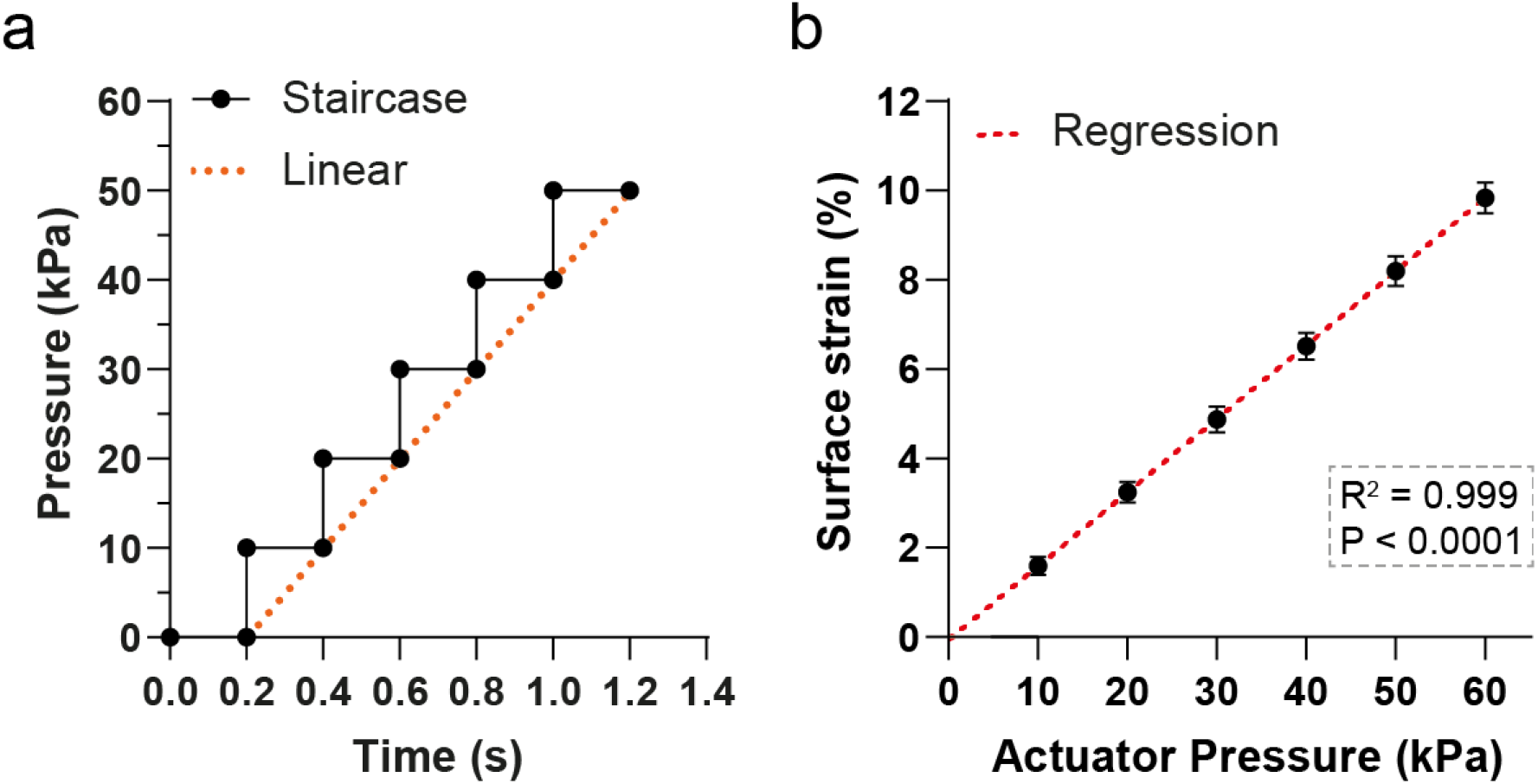
Operation and characterization of the stretching device (**a**) Stretching profile of the pressure actuated stretcher: linear stretch is idealized by a staircase pressure change with adjustable step height and interval. (**b**) Sample calibration showing actuator pressure versus calculated surface strain from the displacement of fiducials markers, (n = 4 membranes, 9-12 different locations/membrane, in total 42.)

